# The contribution of non-additive genetic effects to the genetic variance of polyploid species

**DOI:** 10.64898/2026.05.12.724556

**Authors:** Josselin Clo

**Author notes:** **For correspondence:** Josselin Clo, UMR 8198 – Evo-Eco-Paleo, F-59000 Lille, France. **Social media:** Bluesky @josselin-clo.bsky.social.

## Abstract

Whole genome duplication is a common mutation in eukaryotes with far-reaching phenotypic effects. The resulting morphological, physiological, and fitness consequences and how they affect the survival probability of newly polyploid lineages are intensively studied, but very little is known about the effect of genome doubling on the short-term evolvability of populations. Understanding the effect of polyploidization on the adaptive potential of populations is of crucial importance to predict the future of polyploid populations. In this paper, I investigate the immediate consequences of genome doubling on the genetic variance of populations. To do so, I performed numerical iterations and simulations of how the genetic variance of a quantitative trait changes after polyploidization, under different genetic architectures (additivity, dominance, and epistasis). I found that genetic variance generally decreases after genome doubling. Non-additive gene actions can make autotetraploid populations genetically more diverse than their diploid progenitors in rare cases, notably with overdominance and directional epistasis. By collecting estimates from the agronomic literature, I found that both dominance and epistatic variance contribute to the genetic variance of polyploid populations. These results bring new insights into the adaptive potential of newly formed tetraploid populations, and call for further experimental investigations of how polyploidization is associated with a short-term decrease in evolvability.

## Introduction

Whole genome duplication (WGD hereafter) is common in plants (Ramsey and Schemske, 2002; Soltis *et al*., 2007; Parisod *et al*., 2010; Barker *et al*., 2016) and animals (Mable, 2004), and is considered to have a broad range of effects on plant phenotypes and genomes, and as an important driver of plant adaptation and speciation (Van de Peer *et al*., 2017, 2021). Nevertheless, young autopolyploid lineages, which are the natural outcomes of WGD, are expected to face high extinction rates (see Levin, 2019, for a review). Outside the strong frequency-dependent selection acting on neo-polyploids (i.e. newly formed polyploidy individuals within diploid populations), one historical argument to explain this lower adaptive potential (defined in this manuscript as the amount of genetic variance of populations) of polyploids is that gene duplication dilutes the effects of alleles, such that polyploids are less likely to evolve new adaptive gene complexes compared to diploids (Stebbins, 1971).

How polyploidization can affect the long-term (over hundreds of generations) response to selection and the amount of genetic variance compared to diploids has been an active topic of research, notably in quantitative genetics. For an additive genetic architecture of a quantitative trait, Wright (1938) predicted that, for the same difference in phenotype between homozygous genotypes of diploids and autotetraploids, the additive variance of the tetraploid population is half that of the diploids. When considering the effect of dominance, it is theoretically expected that polyploidization can enhance the genetic diversity of populations in the long term by increasing the frequency of recessive mutations (Haldane, 1927; Ronfort, 1999; Otto and Whitton, 2000; Clo, 2022a, 2022b). The effect of polyploidization on the recombination rate and gametic linkage disequilibrium of a trait also received particular attention (Rowe, 1982; Rowe and Hill, 1984; Griswold and Williamson, 2017), leading to both positive and negative effects pending the genetic architecture of the trait under study. Outside the above-mentioned studies, one underexplored effect of polyploidy on the level of genetic diversity of polyploid populations is the contribution of non-additive genetic effects.

The contribution of non-additive genetic effects to the adaptive potential of populations has mainly been studied in diploid species through the lens of epistasis. It is often assumed that epistasis has a negligible effect on the evolution of populations, because most of the genetic variance estimated in quantitative traits is additive (see Crow, 2010, for a review). However, it has been shown that such studies are misleading, as most of the epistatic interactions affecting the average phenotypic values contribute more to the additive variance than to non-additive ones (Cheverud and Routman, 1995; Hansen, 2013). The consequences of epistatic interactions on breeding values are dependent on whether they tend to decrease or increase, on average, the phenotype of individuals (Carter *et al*., 2005; Le Rouzic *et al*., 2024). In polyploid species, the contribution of non-additive variances can be more important for two reasons. First, polysomic segregation creates the opportunity for more epistatic interactions because of the multiple allele copies per locus (Mostafaee and Griswold, 2019). Second, polysomic inheritance also make that more than one allele per locus to be transmitted from parents to offspring, such that dominance variance contributes partially to the heritable variance of polyploid species (Walsh, 2005).

If there are theoretical expectations about the effect of polyploidization on genetic diversity, empirical patterns are scarcer. Several studies have shown that phenotypic variance can increase after polyploidization (Liu and Wendel, 2003; Pontes *et al*., 2004; Salmon *et al*., 2005). Nevertheless, these studies focused on allopolyploid species, avoiding saying whether the increased variability is due to hybridization or genome doubling *per se*. If we focus on autopolyploids, few estimates of quantitative genetics metrics are available for natural populations. Within those estimates, most of them are estimated in tetraploid populations only, without direct comparison to their diploid progenitors (O’Neil, 1997; Burgess *et al*., 2007), giving limited insights into how autopolyploidization affects the amount of genetic diversity in the short-term (i.e. the very first generations following polyploidization). To my knowledge, two studies experimentally investigated how genome doubling affects the amount of genetic diversity of populations in the short- and/or long-term. Martin and Husband (2012) performed an artificial selection experiment to infer the realized heritability for flowering time in diploid, synthetic-tetraploid, and established tetraploid populations of *Chamerion angustifolium*. They found that newly formed synthetic tetraploids are the most genetically diverse cytotype, while established ones are the worst, diploids being intermediate (Martin and Husband, 2012). Olmedo-Castellanos *et al*. (2025) found that in diploid, tetraploid, and hexaploid populations of *Erysimum incanum*, the ploidy level does not alter the average genetic variance of populations. It appears that few studies focused on the quantitative genetics of autopolyploid species, and that none of these studies tries to decipher the role of additive *versus* non-additive variance on the amount of genetic variance of populations.

Surprisingly, it appears that investigations about the adaptive potential of polyploids are rare in the literature, either theoretically or empirically, which is striking knowing the economic importance of polyploid crops and invasive species. Such knowledge is primordial to understanding the evolutionary dynamics of polyploid species. In this paper, I investigated theoretically how WGD affects the amount of genetic variance of tetraploid lineages when considering non-additive genetic effects. I decided to focus on autotetraploid species to catch the effect of genome doubling *per se*. To do so, I performed numerical iterations and simulations to infer the heritable genetic variance of a quantitative trait subject to different genetic architectures (dominance and epistasis), and I quantified how heritable variance changes in tetraploid populations compared to the diploid progenitors. I found numerically that the genetic variance of a quantitative trait is more likely to decrease after a polyploidization event, independently of the genetic architecture of the trait. By performing a literature survey of agronomic species, I also compared these estimates of the contribution of non-additive variances with empirical data.

## Material and Methods

### Methodological framework and general assumptions

I focused on the quantitative genetics’ definition of evolvability, that is the standardization of the additive variance (*σ*^2^_A-2x_) of a trait by its squared mean *μ*^2^_2x_ (*e*_*2x*_ = *σ*^2^_A-2x_/ *μ*^2^_2x_, Houle, 1992; Hansen *et al*., 2011) for a diploid population, and the standardization of the additive variance and one-third of the dominance variance of a trait by its squared mean (*e*_*4x*_ = (*σ*^2^_A-4x_ + *σ*^2^_D-4x_/3)/ *μ*^2^_4x_, Gallais, 2003 p. 320) for tetraploids. The production of gametes with two alleles results in the inheritance of both additive and dominant genetic effects. That is why the dominance variance also contributes partially to the heritable variance of an autotetraploid population (Walsh, 2005). Evolvability, when quantified in this way, can be interpreted as the proportional change in the trait mean per generation when selection on the trait is as strong as selection on fitness (Hansen *et al*., 2011). Such standardization is needed to compare the adaptive potential among traits (Hansen *et al*., 2011), and it is of particular importance when comparing the evolutionary potential of a trait among cytotypes, because polyploidization is generally associated with an increase in the size of morphological traits (Otto, 2007; Vamosi *et al*., 2007; Porturas *et al*., 2019; Clo and Kolář, 2021). The increase in trait values directly affects the variance of the trait, independently of the other genetic effects associated with polyploidization.

I focused on random mating populations for simplicity, as the decomposition of the genetic variance with inbreeding and dominance is not trivial and necessitates numerous terms (Walsh and Lynch, 2018, chapter 23). Nevertheless, the following results are expected to hold for populations of any selfing rate, since the consequences of genome doubling on genetic variance and the mean of traits are similar in outcrossing and selfing populations. The genotypic frequencies and associated genotypic values at one locus for a bi-allelic model are given in Table 1. I used the Gallais’ (2003, p. 169) nomenclature for simplicity, which assumes that the distance between diploid and tetraploid homozygote genotypes is multiplied by two. However, all the following results are generalizable for any multiplicative factor (see Clo, 2022a; Zwaenepoel, 2025, for examples on additive genetic architectures).

**Table 1.** Genotypic frequencies and values at random mating equilibrium, for diploid and tetraploid populations, for a bi-allelic model (Kempthorne, 1955), *p*_1_ and *p*_2_ are respectively the frequency of the alleles A_1_ and A_2_. The nomenclature of genotypic values is the same than is Gallais (2003, p.169), *a* referring to the distance between homozygote genotypes and their average value, and *d* (respectively *d*_1_, *d*_2_, *d*_3_) the deviation of the heterozygote genotype from additivity in diploids (respectively in tetraploids).

| Diploid population |  |  |  |  |  |
| --- | --- | --- | --- | --- | --- |
| Genotype | $A_1A_1$ | | $A_1A_2$ | | $A_2A_2$ |
| Frequency | $p_1^2$ | | $2p_1p_2$ | | $p_2^2$ |
| Value | $-a$ | | $d$ | | $a$ |
| Tetraploid population |  |  |  |  |  |
| Genotype | $A_1A_1A_1A$ | $A_1A_1A_1A$ | $A_1A_1A_2A$ | $A_1A_2A_2A$ | $A_2A_2A_2A$ |
|  | 1 | 2 | 2 | 2 | 2 |
| Frequency | $p_1^4$ | $4 p_1^3 p_2$ | $6 p_1^2 p_2^2$ | $4 p_1 p_2^3$ | $p_2^4$ |
| Value | $-2a$ | $-a + 3d_1$ | $4d_2$ | $a + 3d_3$ | $2a$ |

I made the hypothesis that the phenotypic values are equal to the genotypic values, such that the environmental effects are null on average, or small enough compared to the genotypic values, such that they are negligible (Falconer and Mackay, 1996, chapter 7).

Finally, I neglected the contribution of other evolutionary forces, such as mutation or selection, in the numerical iterations, because they are not expected to drastically modify the allelic frequencies in the very first generations after genome doubling.

The values of tetraploid genotypes compared to the diploid genotypes, and their relative frequencies, are given in Table 1. Different scenarios of dominance values are given in the supplementary table S1.

### Quantitative genetics of autotetraploid populations

In the first part of this section, I briefly explained how to infer the additive variance in a diploid population and additive as well as dominance variances in an autotetraploid population, as these quantities are necessary for estimating the evolvability of diploid and autotetraploid populations. In a second section, I extended the epistatic model of Cheverud and Routman (1995) for autotetraploid species to study how epistasis modified the evolvability of populations in diploid and autotetraploid populations.

#### Model without epistasis

I considered a single locus with two alleles, A_1_ and A_2_, found respectively at frequencies *p*_1_ and *p*_2_. Let 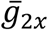 and 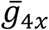 denote the mean genotypic values of the diploid and tetraploid populations at the locus under consideration, calculated as the product of the genotypic values and their frequency (Table 1). The additive variance of a population is the variance of the average effects of alleles at one locus, the average effects of alleles A_1_ and A_2_ being equal to

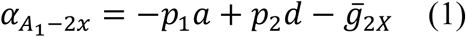

and

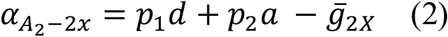

in the diploid population. Similarly, the average effects of alleles in the tetraploid population are

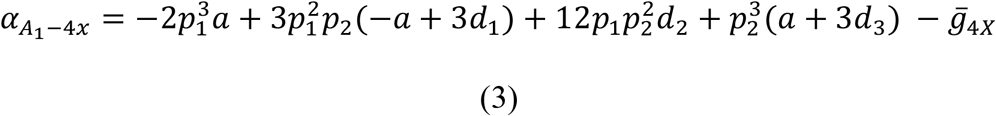

and

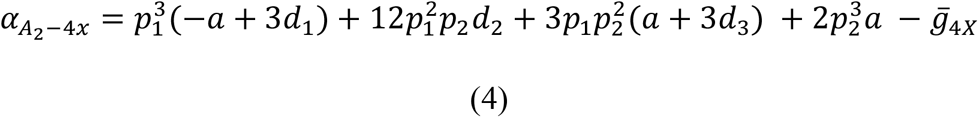

in the tetraploid population (Gallais, 2003, p. 165).

The additive variances of the diploid and tetraploid populations are thus respectively equal to

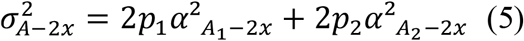

and

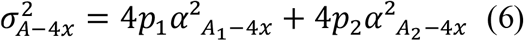

(Gallais, 2003). The production of gametes with two alleles results in the inheritance of both additive and dominant genetic effects. Therefore, dominance variance also contributes to the heritable variance of an autotetraploid population, and it is necessary to infer the first-order (or digenic) interaction terms among alleles. These terms are equal to

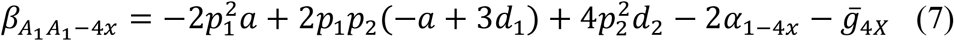

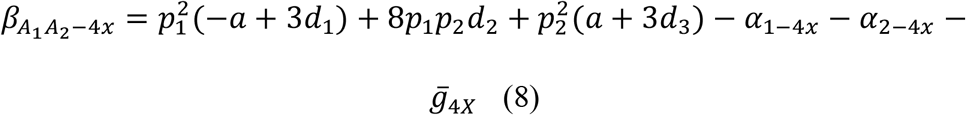

and

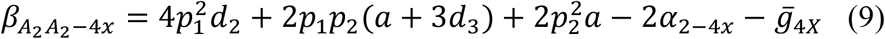

(Gallais, 2003, p. 166). The dominance variance of a tetraploid population is then equal to

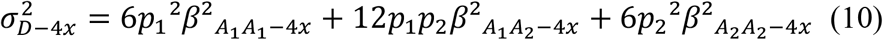

(Gallais, 2003). Different scenario of dominance has been simulated (see Table S1 for details), with partial triplex dominance (at least two A_2_ alleles are needed in heterozygotes to deviate from the mid-homozygote value), duplex dominance (a signle A_1_ allele is needed in heterozygotes to deviate from the mid-homozygote value), and overdominance (all heterozygotes exceed the highest homozygote genotype).

#### Model with epistasis

Epistasis could be a major contributor to a trait’s genetic architecture, since the number of epistatic interactions increases exponentially with polyploidization (Mostafaee and Griswold, 2019). Epistasis occurs when the genotypic value at one locus depends on the genotypic value at another locus. In this section, I extended the Cheverud and Routman (1995) model for autotetraploid species by keeping the nomenclature of Gallais (2003). This parametrization is particularly useful, as it allows us to quantify the contribution of epistasis to the heritable variance. I considered two loci with two alleles, A_1_ and A_2_ at frequency *p*_1_ and *p*_2_ at locus 1, and B_1_ and B_2_ at frequency *q*_1_ and *q*_2_ at locus 2. The subscripts A and B refer to genetic values at locus A and B, respectively (for example, the genotypic values of A1A1 and B1B1 are −2*a*_A_ and −2*a*_B,_ respectively).

The single locus genotypic values are calculated as the unweighted mean of the three or five genotypic values found at the other loci, depending on whether we are considering a diploid or tetraploid population. For example, the single locus genotypic value of the genotypes A_1_A_1_ and A_1_A_1_A_1_A_1_, −*a*_A_ and −*2a*_A_ respectively, are equal to

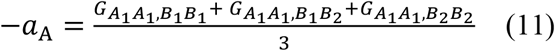

for diploid individuals (Cheverud and Routman, 1995), and

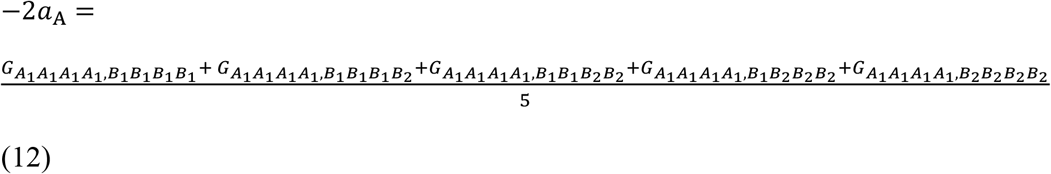

for tetraploids, where *G*_*AXAX*.*BXBX*_ and *G*_*AXAXAXAX*.*BXBXBXBX*_ are the two-locus genotypic values. Similar equations hold for the other genotypes at the first locus and for locus B. Based on this definition, it is possible to infer the two locus non-epistatic values of the genotypes. For example, the non-epistatic values of the genotypes A_1_A_1_-B_1_B_1_ and A_1_A_1_A_1_A_1_-B_1_B_1_B_1_B_1_, *ne*_A1A1,B1B1_ and *ne*_A1A1A1A1,B1B1B1B1_, are equal to

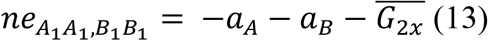

for diploid individuals (Cheverud and Routman, 1995), and

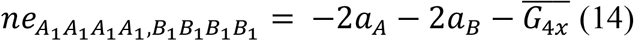

for tetraploid individuals, where 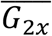 and 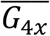 are the unweighted averages of respectively the nine and the 25 possible genotypic values for diploid and tetraploid populations. These non-epistatic genotypic values are the predicted values given by the unweighted least-squares regression of two-locus genotypic values onto the single-locus components (Cheverud and Routman, 1995). The epistatic values of the genotypes are the difference between the two-locus genotypic values and the non-epistatic values. For example, the epistatic values of the genotypes A_1_A_1_-B_1_B_1_ and A_1_A_1_A_1_A_1_-B_1_B_1_B_1_B_1_, *e*_A1A1,B1B1_ and *e*_A1A1A1A1,B1B1B1B1_, are equal to

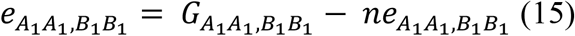

for diploid individuals (Cheverud and Routman, 1995), and

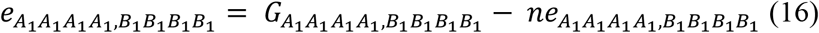

for tetraploid individuals. The mean population values are equal to

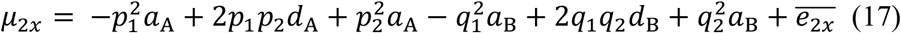

and

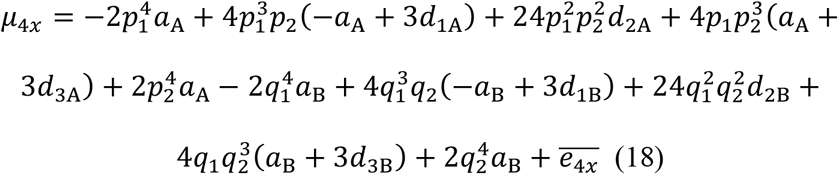

respectively for diploid and tetraploid populations, where 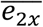 and 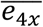 are the populations weighted average epistatic values of the diploid and tetraploid populations (the sum of the product of epistatic values and their frequencies). Based on all these definitions, it is possible to infer the contribution of epistasis into the additive and dominance variances in diploid and tetraploid populations, and the evolvability of each cytotype.

For inferring the single locus components (additive and dominance variance), the procedure is similar to the one presented in the previous section, but we have to take into account that a given genotype transmits its genotypic value (-*a*_A_ or −2*a*_A,_ for example, for the genotype A_1_A_1_ and A_1_A_1_A_1_A_1_) and its associated mean epistatic effect. The mean epistatic effect associated to a given single locus genotype is estimated as

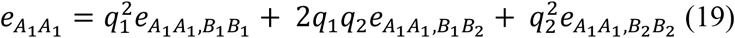

and

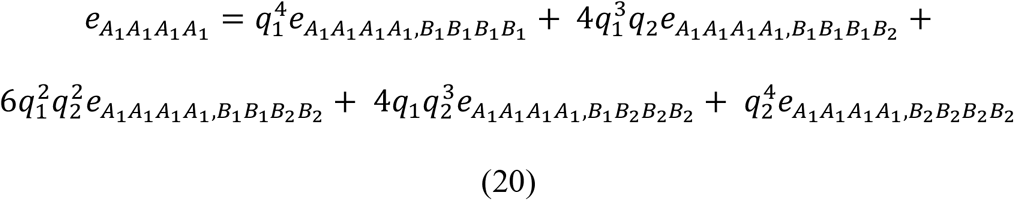

respectively, for the diploid A_1_A_1_ and tetraploid A_1_A_1_A_1_A_1_ genotypes at the first locus. Those equations also hold for the other genotypes at the first locus, and for the genotypes at the second locus. Now, I will define the marginal genotypic (*C*_AXAX_) values as the sum of the genotypic value of a single locus genotype (eqs 11 and 12) and its mean epistatic effect (eqs 19 and 20). For example, the marginal genotypic values of genotypes A_1_A_1_ and A_1_A_1_A_1_A_1_ are equal to

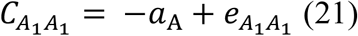

and

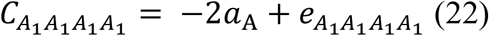

Using the marginal genotypic values in equations 1 to 10 will allow inferring the additive and dominance variances at the two studied loci. One can see that the number of epistatic interactions is much higher in tetraploid populations compared to diploids (nine against twenty-five), and that epistasis contributes to the evolvability of diploids only through the additive variance, while it contributes to tetraploids’ evolvability through both additive and dominance variance. It is also interesting to note that, for the additive scenario, the dominance variance is non-zero because the heterozygote genotypes are not at mid-distance from the homozygote genotypes due to epistasis.

As the sign of epistatic interactions is the key to understanding the effect of epistasis on evolvability (Carter *et al*., 2005), I will test two scenarios, one in which epistasis tends to proportionally increase genotypic values and one in which it tends to proportionally decrease the genotypic values. For simplicity, I will consider additive and fully dominant scenarios, the architecture being the same at the two studied loci. In the model, I will consider epistatic interactions between the allele “A_2_” and “B_2_’. Epistatic effects increase with the number of “A_2_” or “B_2_” copies that a genotype carried, each copy of the above-mentioned alleles increasing or decreasing the genotypic values of 5%. For example, the genotypes A_1_A_1_-B_1_B_1_ and A_1_A_1_A_1_A_1_-B_1_B_1_B_1_B_1_ have unchanged genotypic values, while the genotypes A_2_A_2_-B_2_B_2_ and A_2_A_2_A_2_A_2_-B_2_B_2_B_2_B_2_ respectively have an increase or decrease of genotypic value of 20% and 40%.

#### Review of the decomposition of the genetic variance of agronomic species

I used Google Scholar to perform the literature survey. I used the keywords “polyploid*” and (“genetic variance”, “additive variance”, “dominance variance”, “epistatic variance”), and “plant*”. To be incorporated in the data collection, the selected study had to (1) define clearly the autopolyploidy origin of the species and level of ploidy of the population under study, and (2) describe the amount of additive variance and one non-additive variance (dominance, epistasis, both, or broad-sense heritability). A summary of the sampled species, their ploidy level, and their origin of ploidy can be found in Table S2.

With these estimates, I will infer the difference between the amount of narrow-sense heritability (*h*^2^) and the amount of dominance variance (*h*^2^_D_ = *σ*^2^_D_/*σ*^2^_P_), the amount of epistatic variance (*h*^2^_I_ = *σ*^2^_I_/*σ*^2^_P_), or the broad-sense heritability (*H*^2^ = *σ*^2^_G_/*σ*^2^_P_). I will perform Wilcoxon signed-rank tests to assess if the differences are significantly different from zero.

## Results

### The effect and contribution of dominance to evolvability

With dominance, the effect of polyploidization on evolvability was not constant anymore and depended on the dominance relationships among alleles (Figure 1). It was first notable that, as for the additive scenario, the evolvability of the newly formed tetraploid populations was most of the time shorter than that of the diploid progenitor populations (Figure 1). The decrease was variable and depended on the allele frequency and the dominance level. With partial and complete dominance, the reduction varied from almost 0% to almost 100% (Figure 1A, B, and C). With overdominance, the reduction remained more modest (Figure 1D). The higher the dominance level, the larger the fraction of allele frequency for which autotetraploids were more genetically diverse than diploids, notably for low frequencies of the dominant allele (Figure 1), because of the higher heterozygosity in autotetraploids. When heterozygote genotypes were overdominant, autotetraploids had higher evolvability than diploids. Another interesting result was that, except for the overdominant scenario, the contribution of dominance variance to evolvability was modest in autotetraploid populations (Figure 2).

**Figure 1.**
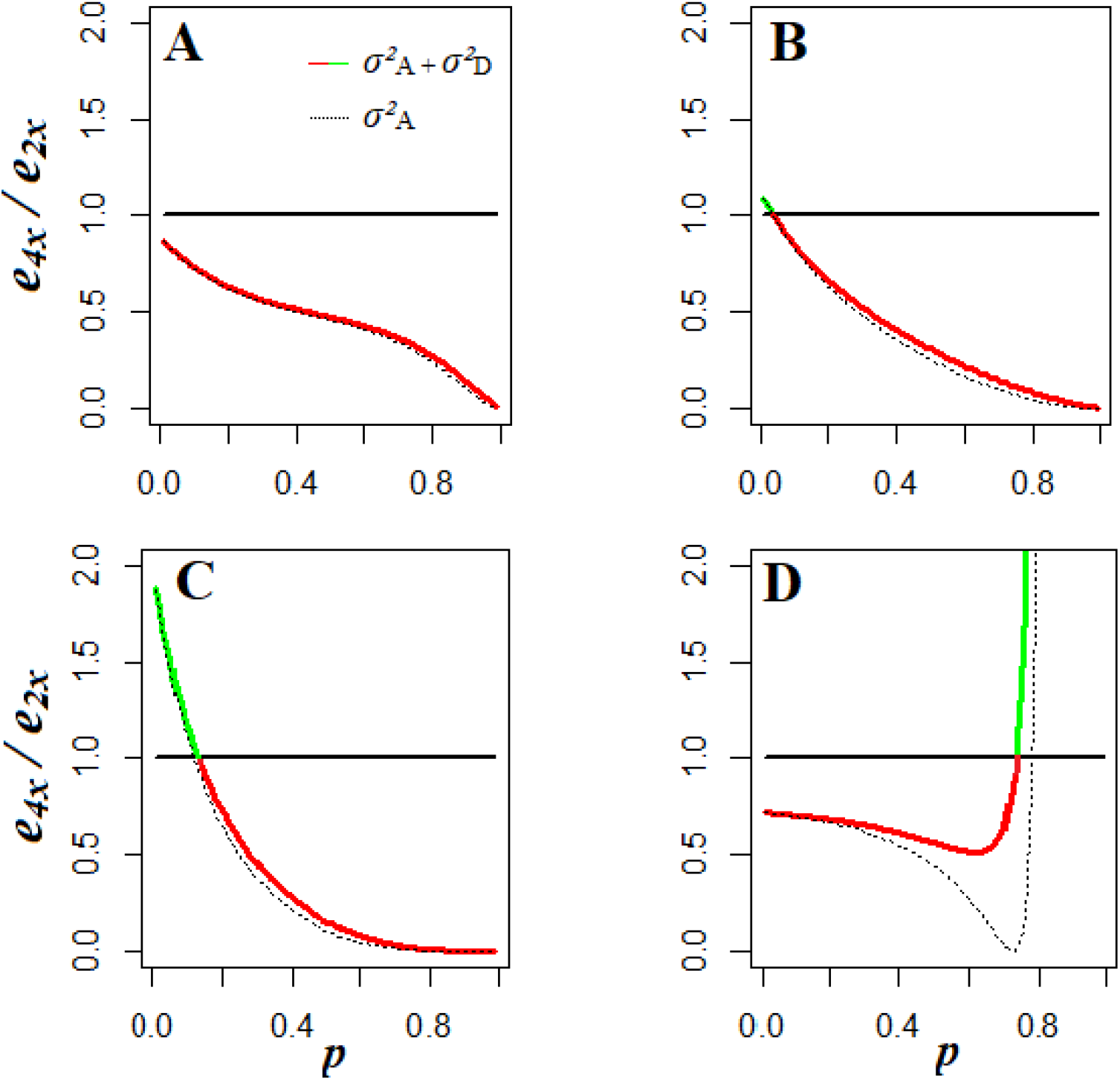
Effects of genome doubling on the evolvability of neo-tetraploids (*e*_4X_), as a function of the allelic frequency of allele A_2_ (*p*_2_), with additive and dominant gene effects. The coloured line represents the evolvability of tetraploids (composed of additive and partly of dominance variance), the line is green when *e*_4x_ > *e*_2x_, and red in the opposite case. the dotted-line represents the partial evolvability of tetraploids (composed of additive variance only). **A.** Partial triplex dominance scenario, with A_1_A_1_A_1_A_1_ = A_1_A_1_A_1_A_2_ > A_1_A_1_A_2_A_2_ > A_1_A_2_A_2_A_2_ > A_2_A_2_A_2_A_2_. **B**. Partial duplex dominance scenario, with A_1_A_1_A_1_A_1_ = A_1_A_1_A_1_A_2_ = A_1_A_1_A_2_A_2_ > A_1_A_2_A_2_A_2_ > A_2_A_2_A_2_A_2_. **C**. Complete dominance scenario, with A_1_A_1_A_1_A_1_ = A_1_A_1_A_1_A_2_ = A_1_A_1_A_2_A_2_ = A_1_A_2_A_2_A_2_ > A_2_A_2_A_2_A_2_. **D**. Overdominant scenario, with A_1_A_1_A_1_A_2_ = A_1_A_1_A_2_A_2_ = A_1_A_2_A_2_A_2_ > A_1_A_1_A_1_A_1_ > A_2_A_2_A_2_A_2_.

**Figure 2.**
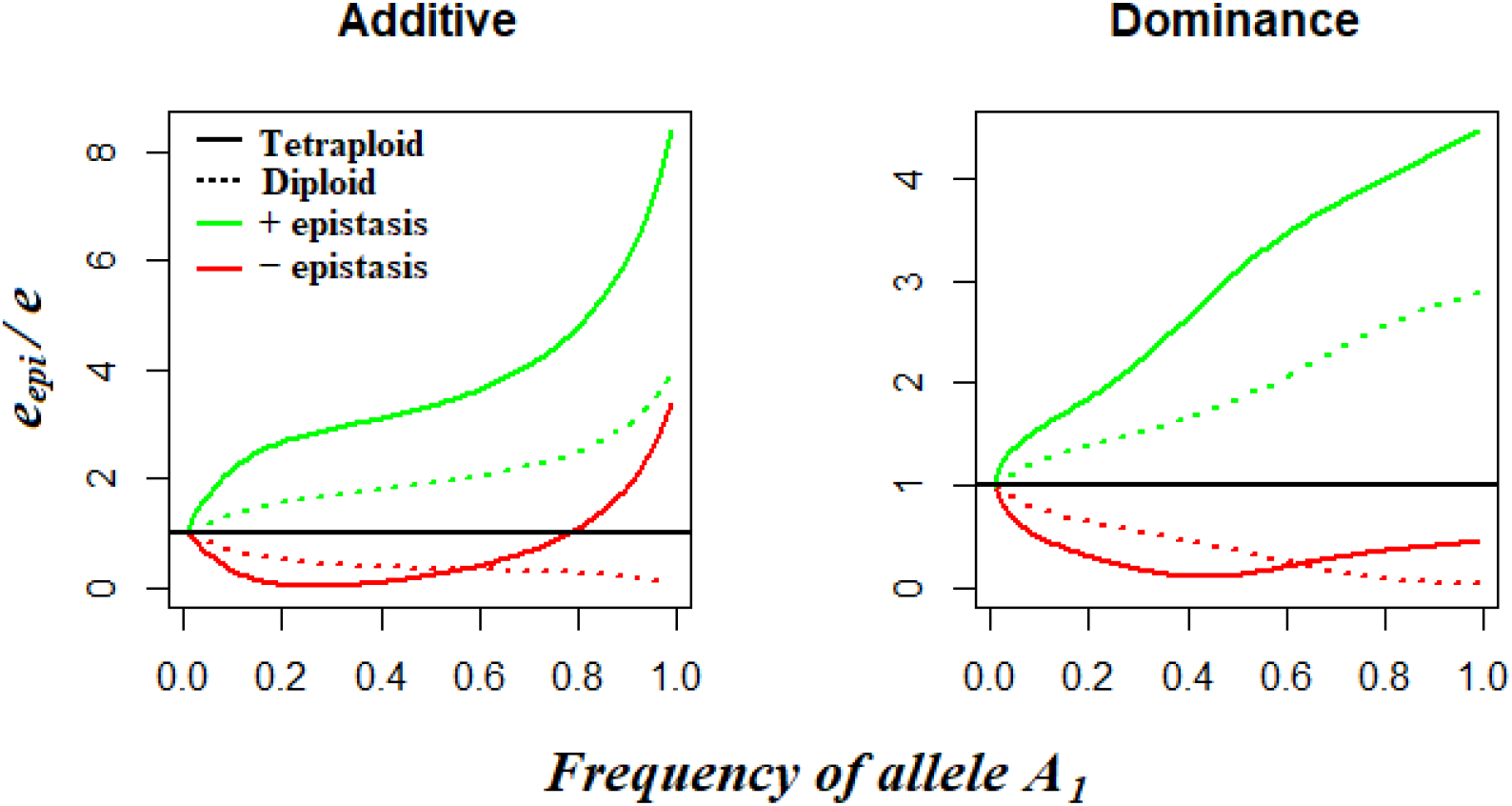
Ratio of evolvability with epistasis over the evolvability without epistasis (*e*_*epi*_/*e*) as a function of the frequency of allele A_2_, for tetraploid and diploid populations (respectively solid and dotted lines) and for positive and negative epistasis (respectively green and red lines), for the additive and dominant genetic architecture of the quantitative traits. The frequency of allele B_2_ is fixed and equal to 0.25. The black horizontal line represents the threshold for which evolvability with epistasis is equal to evolvability without epistasis.

As for the additive scenario, when the founding population was small and stochasticity high, the decrease in evolvability of autotetraploids was higher than expected in numerical iterations (Figure S1).

### Does epistasis affect diploids and tetraploids equally?

This section studied how the different forms of epistasis (positive or negative) affect the evolvability of each cytotype, separately. For clarity, I presented 2D graphics in the main text, by plotting how evolvability changed as a function of the frequency of allele A_1_, the frequency of allele B_1_ was fixed and equal to 0.25. The heatmaps presenting the evolution of evolvability as a function of the frequencies of A_1_ and B_1_ alleles can be found in the supplementary materials (Figures S2 and S3).

With or without dominance, the qualitative effects of epistasis on evolvability within cytotypes were the same, with positive and negative epistasis respectively increasing and decreasing evolvability, on average, for both cytotypes (Figures 2, S2, and S3). It was nevertheless remarkable that for tetraploids, positive epistasis led to a stronger increase in evolvability compared to diploids (Figures 2, S2, and S3). Negative epistasis was less often negative for evolvability in tetraploid populations (Figures 2, S2, and S3), and often had a smaller effect on evolvability in tetraploids than in diploids (Figures 2, S2, and S3). For negative epistasis in the additive scenario, negative epistasis can even increase the evolvability of tetraploids compared to a scenario without epistasis (Figure 2). This is because, with epistasis, the genetic architecture of the trait was not additive anymore, and dominance relationships among alleles within a locus appeared. With negative epistasis, the alleles A1 and B1 became recessive. Recessive actions of genes had a symmetrical effect, it increased the evolvability of tetraploid populations compared to an additive scenario, but for high frequencies (Figure 2).

### For which allelic frequencies does epistasis increase the evolvability of tetraploids?

This section studied the conditions under which epistasis can make tetraploids more genetically diverse than their diploid progenitors. For clarity, I present heatmap graphics in the main text, by plotting how evolvability changed as a function of the frequency of the alleles A_1_ and B_1_ and epistasis, with the upper-limits of ratios of evolvability being fixed at 2. The untransformed heatmaps can be found in the supplementary materials (Figure S4).

The main effect of positive epistasis was to increase the value of the ratio of tetraploid evolvability over diploid evolvability, in both additive and dominant scenarios (Figure 3), compared to non-epistatic scenarios (Figure 2). Nevertheless, the greater gain in evolvability due to epistasis for tetraploids (Figure 2) was most of the time not enough to overwhelm the loss of evolvability associated with genome doubling (Figures 3 and S4), with most of the frequencies representing points for which *e*_4x_ are smaller than *e*_2x_. Independent of the sign of epistasis, scenarios for which polyploids have higher evolvability than diploids are more often reached when one of two epistatic alleles is found at high frequency and the other at low frequency (Figure 3 and S4). Negative epistasis had opposite effects, either decreasing or increasing the ratio of evolvabilities, depending on allelic frequencies (Figures 3 and S4). Notably, negative epistasis can make tetraploids more genetically diverse than diploids (Figures 3 and S4), because negative epistasis has a more drastic effect in diploids when the allele A_1_ is in high frequency in populations (Figure 2). Positive epistasis does not affect the prediction that, on average, diploids had higher evolvability than tetraploids, despite the stronger effect of epistasis expected in polyploids (Figure 3 and S4).

**Figure 3.**
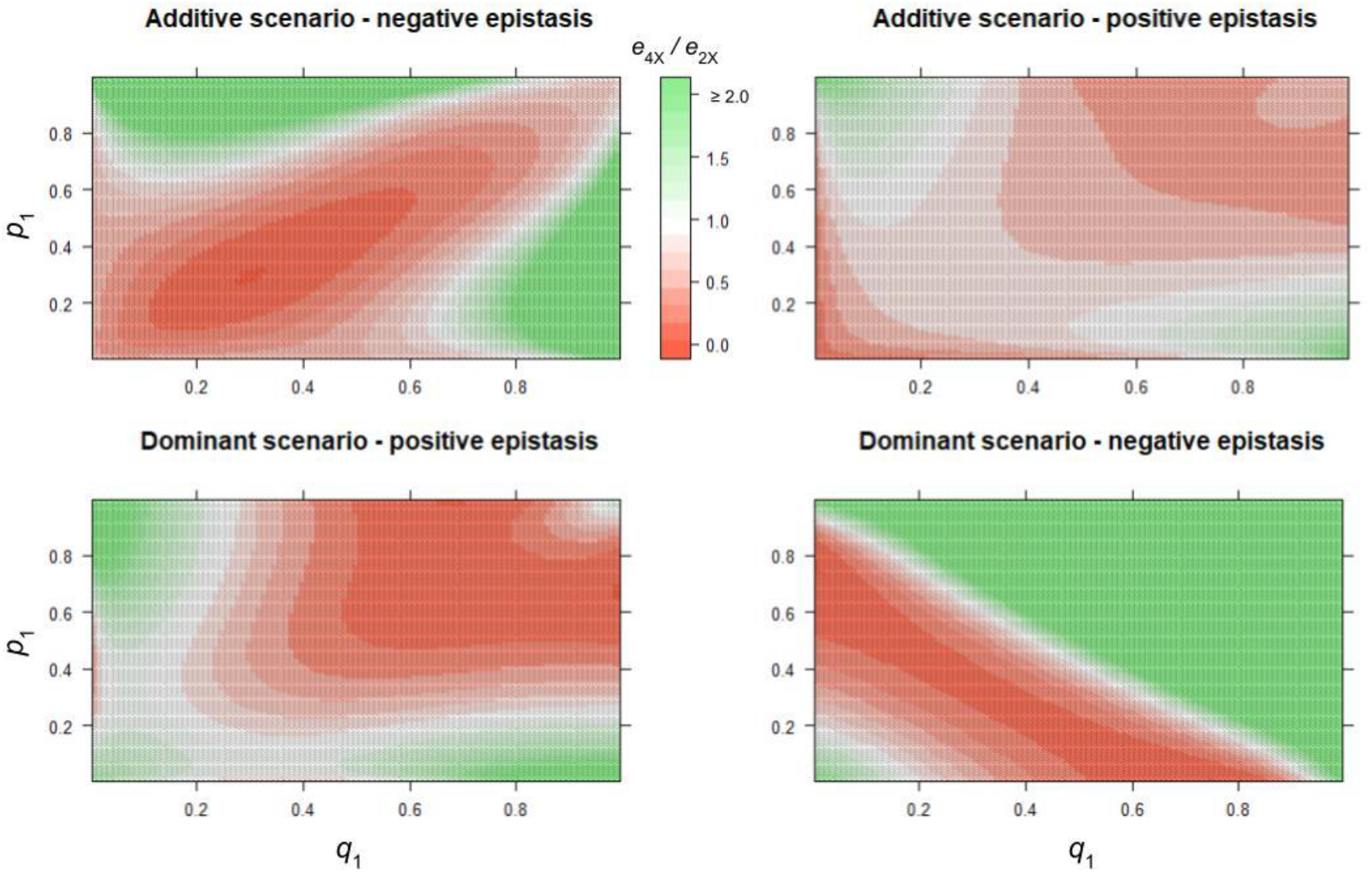
Heatmap of the effects of genome doubling and epistasis on the ratio evolvability of neo-tetraploids (*e*_4X_) over diploids (*e*_2X_), as a function of the allelic frequency of allele A_2_ (*p*_2_) and B_2_ (*q*_2_), with additive and dominant gene effects in the presence of epistasis (positive or negative). The heatmap is green when *e*_4x_ > *e*_2x_, and red in the opposite case. For clarity, the upper-limit of the ratio is set to 2, but it can be much bigger (see supplementary material S4 and S5 for full details).

#### Contribution of non-additive variance in polyploid plant species

In total, I sampled 78 estimates, all found for agronomic autopolyploid species, for which the quantitative genetics of polyploid species is much ahead. Those estimates are divided among 12 species (Table S2).

From the comparisons of additive variance with non-additive variances, it seems that, for autopolyploid plant species, dominance and epistatic variances contribute to the same order of magnitude as additive variance (Table 2, Figure 4) when taken alone, but that most of the total phenotypic variance is non-additive when comparing narrow- and broad-sense heritabilities (Table 2, Figure 4). The variation of estimates is, however, important (Figure 4).

**Table 2.** Comparison of additive variance with non-additive genetic diversity (± standard deviation) in autopolyploid agronomic plant species. Further details are given in the supplementary Table S2.

| Comparison | *N* | Estimate (± s.d) | *p*-value |
| --- | --- | --- | --- |
| $h^2 - h^2_{\text{D}}$ | 65 | -0.050 (0.513) | 0.597 |
| $h^2 - h^2_{\text{I}}$ | 34 | -0.031 (0.277) | 0.746 |
| $h^2 - H^2$ | 58 | -0.383 (0.288) | < 0.0001 |

**Figure 4.**
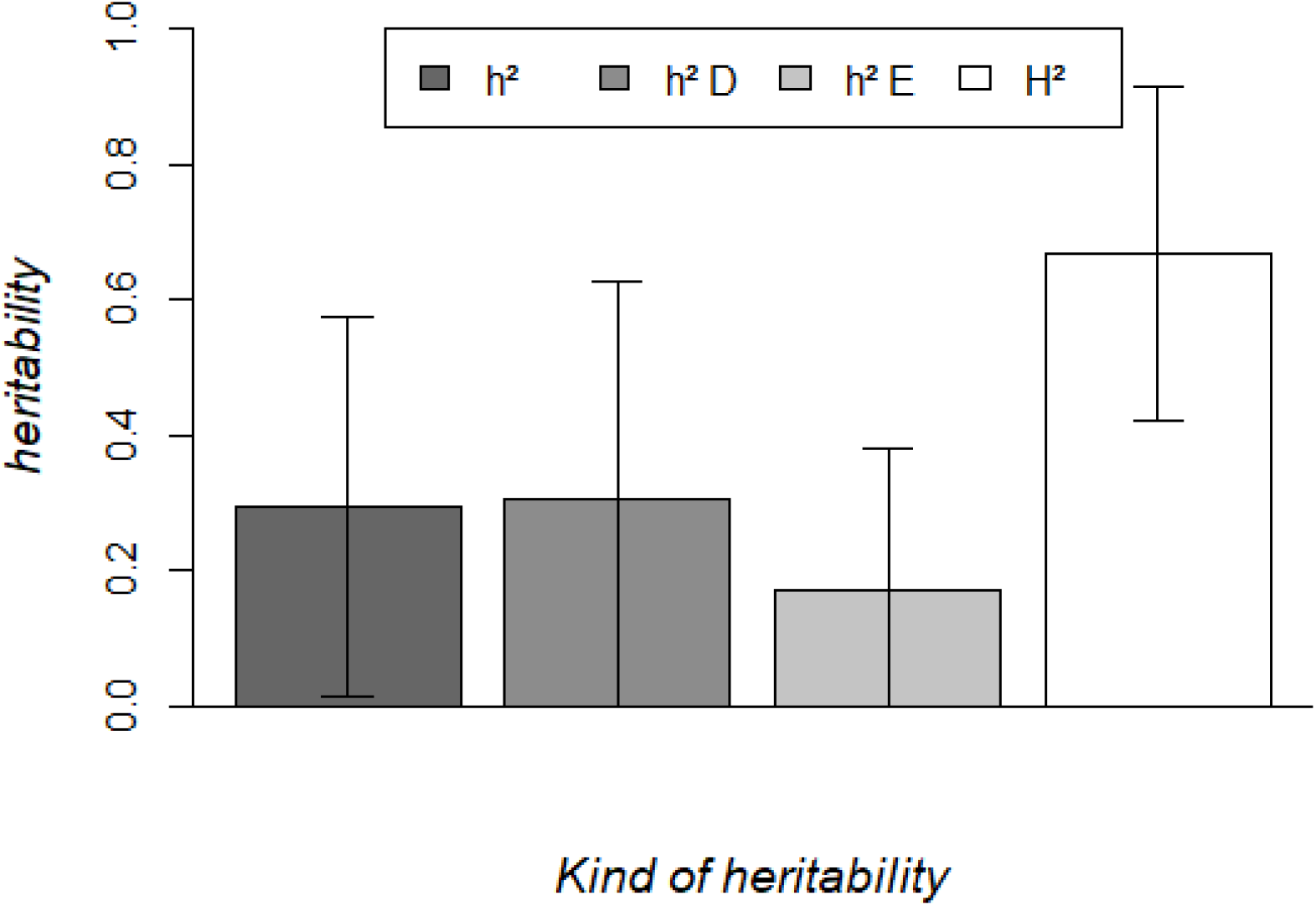
Different heritabilities (+/− standard deviation) for different traits from the agronomic literature of autopolyploid species. h^2^ = narrow-sense heritability (additive variance / phenotypic variance). h^2^ D = dominance variance/ phenotypic variance. h^2^ E = epistatic variance/ phenotypic variance. H^2^ = broad-sense heritability (genetic variance / phenotypic variance).

## DISCUSSION

In this paper, I investigated the effect of whole genome duplication on the evolvability, defined as the heritable variance of a trait standardized by its square mean, of the newly formed tetraploid population. I found that polyploidization is generally deleterious for the amount of genetic variance of populations, independently of the genetic architecture of the modelled trait. Non-additive gene actions can make autotetraploid populations to have higher evolvability values than their diploid progenitors in rare cases, notably with overdominance and directional epistasis.

### Short-term genetic variance of autotetraploid populations

My results have shown that evolvability generally decreases in the very first generations for polyploid lineages, even if dominance and epistasis can contribute to making newly formed tetraploids more genetically diverse than their diploid progenitors. Interestingly, the only study comparing the amount of heritable variance of synthetic neo-tetraploids with their diploid progenitor found that newly formedtetraploids of *Chamerion angustifolium* have the highest realized heritability (Martin and Husband, 2012). This surprising result remains unresolved, even if the authors have proposed explanations. As shown before, the genetic architecture of the trait has important effects on the extent of genetic variance, with dominance and epistasis being able to make autotetraploids more genetically diverse than their diploid progenitors. Interestingly, for quantitative traits under strong stabilizing selection, frequencies of the newly favourable alleles are low (<0.2, Clo and Opedal, 2021). Under these conditions, non-additive effects are expected to increase the evolvability of tetraploid populations, compared to their diploid progenitors (Figures 1 & 3). Dominance and epistasis are known to contribute to the genetic architecture of quantitative traits (Monnahan and Kelly, 2015; Oakley *et al*., 2015; Clo *et al*., 2021; Le Rouzic *et al*., 2024). Notably, epistatic interactions are expected to increase exponentially with polyploidization events (Bingham *et al*., 1994; Griswold and Williamson, 2017), and are theoretically known to play a role in autotetraploid population adaptation (Mostafaee and Griswold, 2019). I only considered a two-locus model in this paper, but the results are expected to be generalized to a higher number of loci (they should sum up), both for dominant and epistatic architecture (if epistasis remains directional). Such an effect of epistasis could be particularly important in selfing species, in which large numbers of genetic interactions are maintained due to the reduced effective recombination rate of populations (Lande and Porcher, 2015; Abu Awad and Roze, 2018; Clo *et al*., 2020; Clo and Opedal, 2021). It would be of great interest to empirically compare the genetic architecture of quantitative traits among diploids and autotetraploids to understand if non-additive genetic effects can be major drivers of autotetraploid genetic variance.

### Autopolyploidization as a source of evolutionary novelties

In addition to modifying the capacity of populations to respond to natural selection, autopolyploidization also leads to major morphological and physiological transformations, some of those potentially affecting the survival probability of newly formedtetraploid lineages. Polyploidization is associated in the short-term with an increase in the size of morphological and cellular traits, and a reduction of fitness (Otto, 2007; Vamosi *et al*., 2007; Porturas *et al*., 2019; Clo and Kolář, 2021). Cellular and fitness modifications seem to be obstacles to the establishment in newly formed polyploid lines, as these components return to diploid-like values (Clo and Kolář, 2021) after establishment, but the increase of morphological structures remain observable in established lineages and comparable to what is found in newly formed polyploids (Clo and Kolář, 2021). This suggests that those modifications could have a role in the survival probability of lineages (Bomblies, 2020), even if these results have been obtained on a restricted number of studies, and that more examples are needed to confirm these patterns. Modifications of morphological traits can theoretically increase the probability of survival of new lineages, if the population undergoes an environmental change and if this change is aligned with the phenotypic modifications in tetraploid individuals (Oswald and Nuismer, 2011; Cheng et al., 2026). As an empirical support, an increase in the size of autotetraploids is known to interact with the local environment to increase the interspecific competitiveness of tetraploids compared to diploids (Čertner *et al*., 2019). In addition to the potential increase in competitive capacities, polyploid lineages also present increased tolerance to stressful conditions compared to diploid populations (see Bomblies, 2020 for a review), with, for example, increased tolerance to drought and salinity (Yang *et al*., 2014; Chen *et al*., 2020). But again, those examples are obtained from very specific stress on a few sets of synthetic polyploids, such that more studies are needed to understand how this can be generalized to other stress and natural polyploids. These different short-term advantages can counteract the initial decrease in genetic diversity predicted in newly formed tetraploid populations by increasing the survival probability of neo-tetraploid populations.

### What do we know from genetic diversity in established tetraploids?

If empirical investigations of the short-term consequences of genome doubling are lacking in the literature, little is known about the long-term consequences of autopolyploidization. Martin and Husband (2012), by using an artificial selection experiment, found that the realized heritability for flowering time is 0.31 in established autotetraploids of *Chamerion angustifolium*, while diploid populations of the same species exhibited a significantly higher realized heritability of 0.40 in the same experiment. Olmedo-Castellanos et al (2025) found that autotetraploid and autohexaploid populations of *E. incanum* were as genetically diverse as their diploid counterparts. No other direct comparisons among autotetraploids and diploids of the same species in controlled environments are available in the literature, but the few estimates of heritability found in natural autotetraploid populations are in the range of what is found in diploid populations (O’Neil, 1997; Burgess *et al*., 2007; Clo *et al*., 2019).

With this very limited set of studies, it appears that established autotetraploids do not more genetic variance than diploids species, despite the less efficient purging of recessive deleterious mutations (Ronfort, 1999; Otto and Whitton, 2000), and the contribution of dominance and epistatic variance in the heritability of autopolyploid lineages I detected in agronomic polyploid plant species (Tables 2 and S2, Figure 4). Interestingly, it seems from the small sample of species examined here, that the contribution of non-additive variances is more important in autopolyploid species than in diploid ones (see, for example, Wolak & Keller 2014 for a comparison with domesticated diploid species). Nevertheless, more estimates are necessary to have a precise idea of how polyploidization affects evolvability in the long term. The discrepancy between the high level of non-additive variance in the data can have multiple origins. First, in the theoretical model, we only focused on digenic dominance variance, while empirical estimates merge all forms of dominance (digenic, trigenic…). The same applies to the different forms of epistasis. Second, we only modeled some scenarios of dominance and epistasis; it is possible that other genetic architectures would lead to different results in the theoretical models. Artificial selection experiments seem the most straightforward way to accurately study this question, due to the contribution of dominance variance to adaptive potential. Those experiments allow to catch the realized heritability or evolvability of a population (containing both additive and dominance contributions). Decomposing the genetic variance of a populations from sibling or pedigree analyses, and then inferred the evolvability could lead to biased estimates, notably due to the limited statistical power to infer correctly the dominance variance in such experiments (Wolak and Keller, 2014; Walsh and Lynch, 2018).

## Conclusions

In this paper, I theoretically investigated how genome doubling affects the genetic diversity of newly formed autotetraploid populations. I found that, in general, the evolvability of newly formed tetraploids decreases in the first generations compared to diploid populations, even if dominance and epistasis can make tetraploids to have more genetic variance than their diploid progenitors. I also found that those non-additive sources of diversity contribute as importantly as additive variance to the total phenotypic diversity. These results introduce a new hypothesis that helps to understand the high extinction rate associated with young polyploid lineages (Levin, 2019): they suffer from an initial decrease in genetic variance, reducing their capacity to cope with new or changing environments. Now, experimental investigations of this hypothesis are needed, in order to understand how and when neo-tetraploid populations are more likely to extinct due to a lack of genetic variance. Artificial selection experiments involving diploid, synthetic tetraploid, and potentially established tetraploid populations seem ideal to go infer the realized heritability/evolvability of quantitative traits among cytotypes (Martin and Husband, 2012). Such experiments will help understanding how genome doubling modifies the genetic diversity of a population in both short- and long-term.

## Acknowledgment

I thank Filip Kolář for comments on the early version of the manuscript. I thank the associate editor and the two anonymous reviewers for their helpful comments, which significantly improved the clarity and quality of the manuscript.

## Data availability

The code and empirical data used in the manuscript can be found at: https://github.com/JosselinCLO/evol_polyploidy_dom_epi.

