## Supplementary figures and tables for "The contribution of non-additive genetic effects to the genetic variance of polyploid species"

**Supplementary materials:**

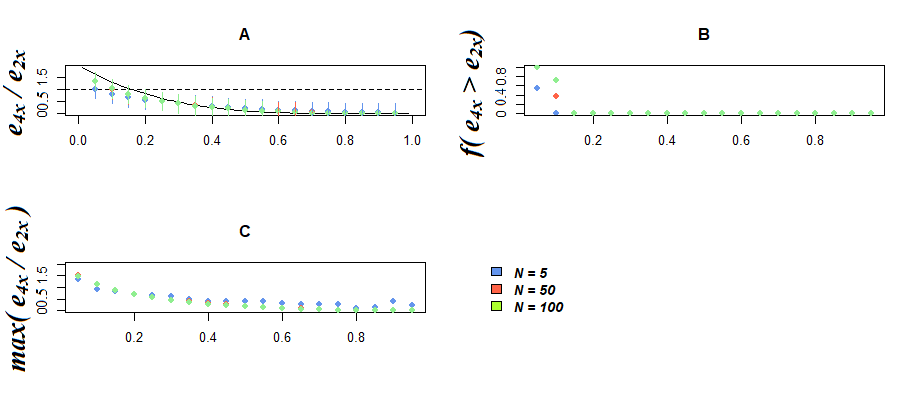

**Figure S1.** Effects of genome doubling on the evolvability of neo-tetraploids (*e*_4X_), as a function of the allelic frequency of allele A_1_ (*p*), in a complete dominance scenario, with A_1_A_1_A_1_A_1_ = A_1_A_1_A_1_A_2_ = A_1_A_1_A_2_A_2_ = A_1_A_2_A_2_A_2_ > A_2_A_2_A_2_A_2_. **A.** Ratio of the evolvabilities of neo-tetraploids (*e*_2X_) over diploid values (*e*_2X_), the solid line is the analytical prediction, color points are simulations results for different size of funding populations, errors bars stand for 95% confidence intervals (*n* = 1000). **B.** Proportion of simulation in which the evolvability of neo-tetraploids is strictly higher than the evolvability of their diploid progenitors, for different size of funding populations. **C.** Maximum value of the ratio of the evolvabilities of neo-tetraploids over diploid values within 1000 simulated populations, for different sizes of funding populations.

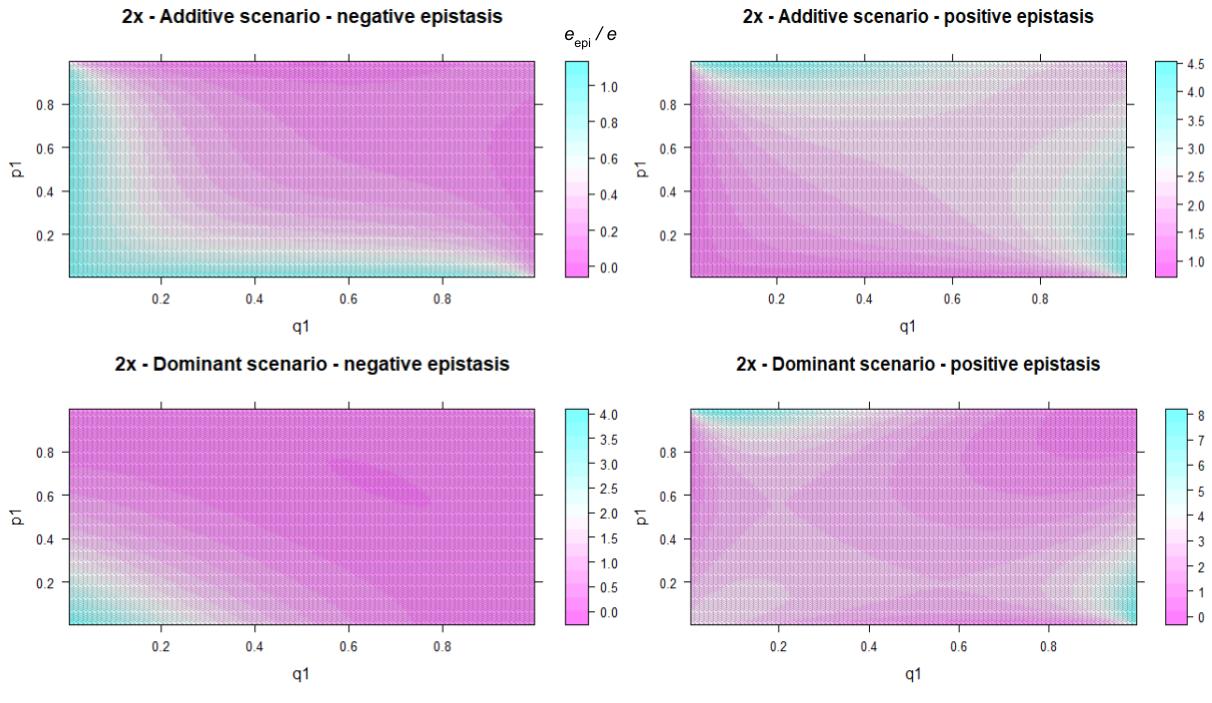

**Figure S2.** Heatmap of the effect of epistasis on the evolvability of diploids, as a function of the allelic frequency of allele A_1_ (*p*_1_) and B_1_ (*q*_1_), with additive and dominant gene effects in presence of epistasis (positive, or negative).

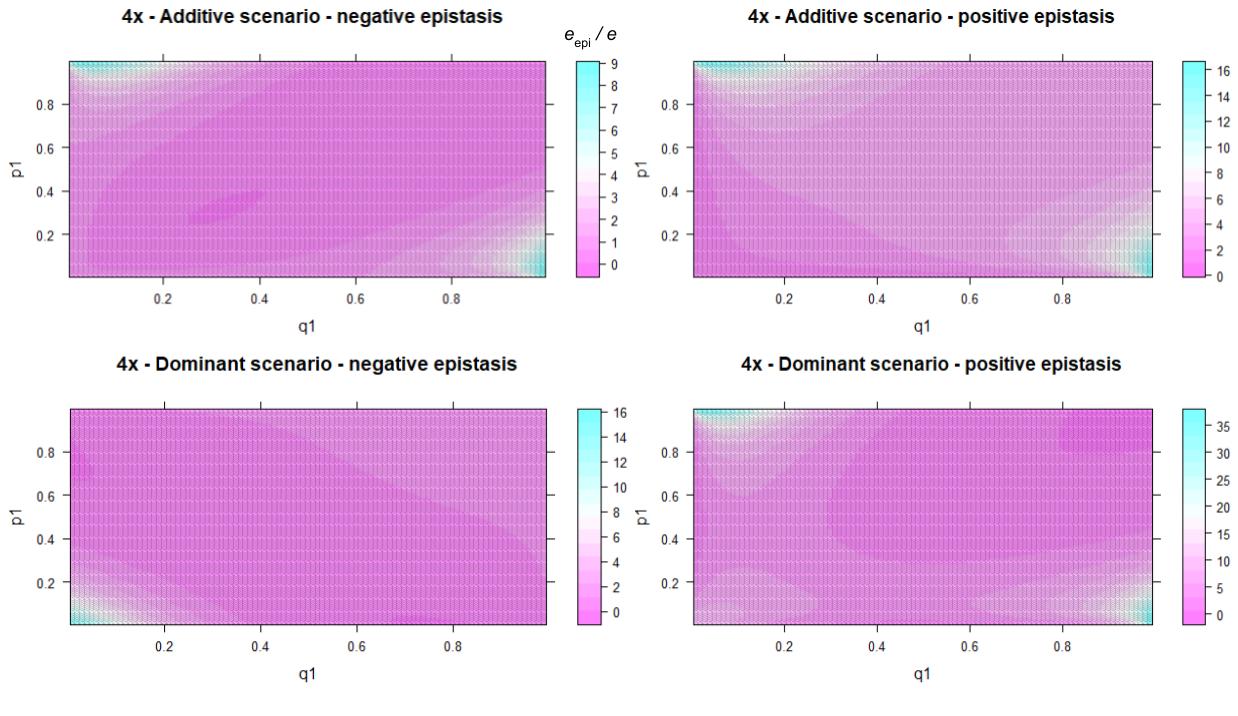

**Figure S3.** Heatmap of the effect of epistasis on the evolvability of neo-tetraploids, as a function of the allelic frequency of allele A_1_ (*p*_1_) and B_1_ (*q*_1_), with additive and dominant gene effects in presence of epistasis (positive, or negative).

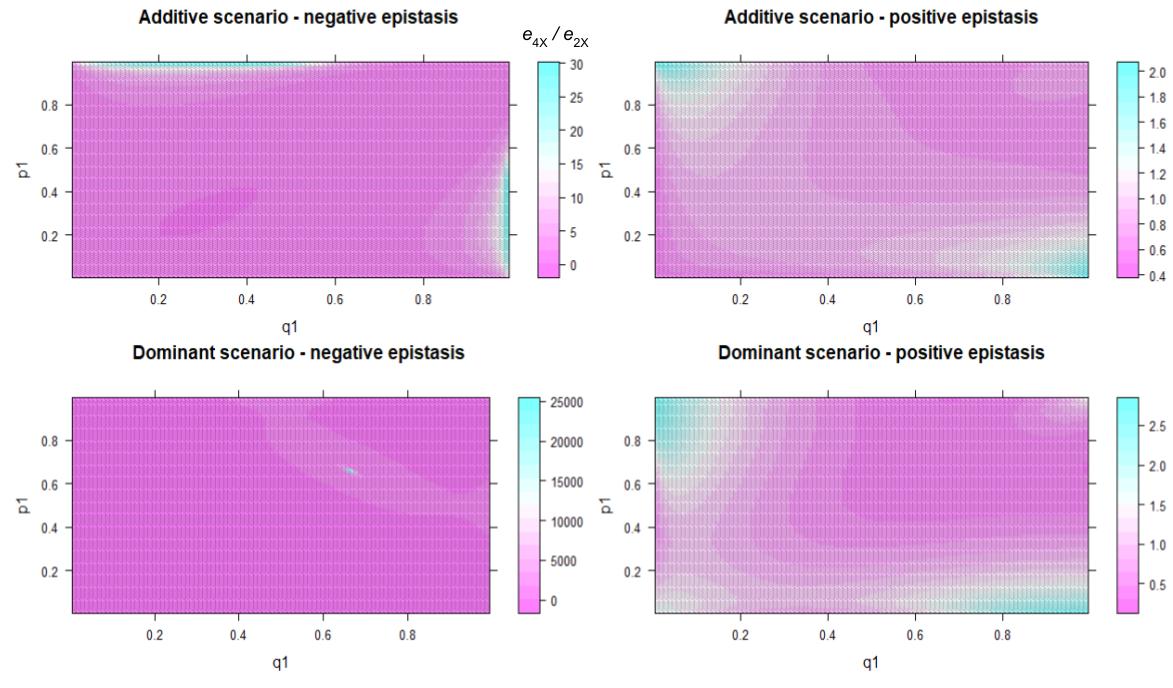

**Figure S4.** Heatmap of the effects of genome doubling and epistasis on the ratio evolvability of neo-tetraploids (*e*_4X_) over diploids (*e*_2X_), as a function of the allelic frequency of allele A_1_ (*p*_1_) and B_1_ (*q*_1_), with additive and dominant gene effects in presence of epistasis (positive, or negative).

**Table S1.** Values of the tetraploid genotypes as a function of the diploid values, for the different genetic architectures simulated.

| **Scenario** | **A_1_A_1_A_1_A_1_** | **A_1_A_1_A_1_A_2_** | **A_1_A_1_A_2_A_2_** | **A_1_A_2_A_2_A_2_** | **A_2_A_2_A_2_A_2_** |
| --- | --- | --- | --- | --- | --- |
| Additive | -2*a* | $-a$ | 0 | $a$ | 2*a* |
| Partial triplex dominance | -2*a* | 0 | 1.5*a* | $2a$ | 2*a* |
| Partial duplex dominance | -2*a* | *a* | 1.5*a* | $2a$ | 2*a* |
| Complete dominance | -2*a* | 2*a* | 2*a* | 2*a* | 2*a* |
| Overdominant | -2*a* | 2.2*a* | 2.2*a* | 2.2*a* | 2*a* |

**Table S2.** List of species sampled for the literature survey with the number of observations, the ploidy level, and their ploidy origin.

| Species | Number of observations | Ploidy level | Ploidy origin | References |
| --- | --- | --- | --- | --- |
| *Vaccinum corymbosum* | 10 | Tetraploid | Autopolyploid or segmental allopolyploid with tetrasomic inheritance | Amadeu et al. (2020) |
| *Solanum tuberosum* | 14 | Tetraploid | Autopolyploid | De Bem Oliveira et al. (2019); Amadeu et al. (2020); Panday et al. (2022); Endelman (2023) |
| *Oryza sativa* | 29 | Tetraploid | Autopolyploid | Shadid et al. (2011); Shadid et al. (2012) |
| *Urochloa spp.* | 12 | Tetraploid | Autopolyploid | Matias et al. (2018) |
| *Panicum maximum* | 6 | Tetraploid | Autopolyploid | Lara et al. (2019) |
| *Musa spp.* | 7 | Triploid | Autopolyploid | Tenkouano et al. (2012) |
